# Loss of Aurora kinase signaling allows lung cancer cells to adopt an alternative cell cycle and form multinucleated polyploid giant cells that resist anti-mitotic drugs

**DOI:** 10.1101/865337

**Authors:** Vural Tagal, Michael G. Roth

## Abstract

Polyploid multinucleated (PP/MN) giant cancer cells are common in tumors and have been associated with resistance to cancer therapy, tumor relapse, malignancy, immunosuppression, metastasis, cancer stem cell production and modulation of the tumor microenvironment. However, the molecular mechanisms that cause these cells to form are yet known. In this study, we discover that Aurora kinases are synergistic determinants of a switch from the proliferative cell cycle to polyploid growth and multinucleation in lung cancer cell lines. When Aurora kinases are inhibited together, lung cancer cells uniformly grow into PP/MN giant cells. These cells have adopted an endocycle in which the genome replicates, mitosis is omitted and cells grow in size. Consequently, such cells continue to safely grow in the presence of antimitotic agents. These polyploid multinucleated cancer cells can re-enter the proliferative cell cycle and grow in cell number when the treatment is terminated.

## Main

The goal of cancer medicine is to eradicate tumors entirely. However, a major challenge in treating cancer is posed by intratumor heterogeneity^1,2^. The body of a tumor includes a diverse collection of cells harboring distinct molecular profiles with differential levels of sensitivity to treatment. The tumor population as a whole has the potential to access all the cellular mechanisms encoded in the genome and drug treatment selects for those cells that happen to have adopted an escape mechanism. To combat this, a goal of cancer medicine is to better understand the compositional and functional complexity of tumors in order to find more effective strategies to overcome the disease.

A well-known cancer cell type in tumor populations is called polyploid multinucleated (PP/MN) giant cancer cells ^3–5^. These cells contain multiple nuclei with significantly elevated genomic content (sometimes more than 100 fold) when compared to other cancer cells within the same tumor. The frequency of giant cells can increase markedly in high-grade malignant cancers or following anticancer treatment. They can remain viable and metabolically active for extended periods of time. They can secrete an array of growth factors, cytokines and chemokines ^3,6^. Immunosuppressive proteins including programmed death-ligand 1 (PD-L1) were also found to be overexpressed in these cells ^7^. Therefore, they are responsible of contributing to a microenvironment advantageous for tumor growth and survival.

PP/MN cancer cells can repopulate, and form macroscopic colonies and spheroids in vitro as they generate tumors when inoculated into mice ^8–10^. These cells gain the properties of cancer stem cells and produce daughter cells through asymmetric cell division patterns ^11–13^. Compared with diploid cancer cells, these daughter cells express fewer epithelial markers and acquire a mesenchymal phenotype, which is a key transformation for cancer development, progression and metastasis ^10,14,15^.

Commonly, the ratio of PP/MN cancer cells is higher in high-grade malignant tumors than in low-grade tumors, in relapsing tumors after chemotherapy than in tumors before chemotherapy, and in the metastatic foci than in the primary tumor ^16^. Such cells are known to escape from cytotoxicity induced by major anti-mitotic agents including taxanes and vinca alkaloids ^17–21^. Emerging evidence has demonstrated that PP/MN cancer cells arise in lung cancer ^20^, cervical carcinoma ^22^, ovarian cancer ^10,23^, prostate cancer ^24,25^, gliobastoma ^26^, colorectal cancer ^11,16,27^, and breast cancer ^15,28,29^. The role of PP/MN giant cancer cells and their daughter cells in resistance to therapy, tumor relapse, metastasis, immunosuppression, modulation of the tumor microenvironment and generation of cancer stem cells is well documented. Therefore, revealing the molecular events that cause PP/MN giant cells to form can lead to clinically relevant approaches for treating recurrent and metastatic disease, which remains a major global health issue despite extensive research over the past century. In the course of studying hypersensitivity to inhibitors of Aurora kinase A ^30^, we previously reported an unusual dose response relationship to Aurora kinase inhibitors (AURKi). In some non-small cell lung cancer (NSCLC) lines, more cells survived the inhibitor treatment and kept growing at high than at lower drug concentrations. We show in the work presented below that many NSCLC cell lines grow and uniformly form PP/MN giant cancer cells by adopting an endocycle when Aurora kinases are inhibited. Endocycles omit mitosis, do not require the mitotic machinery and therefore, provide a safe path for growth by avoiding the catastrophic consequences of anti-mitotic perturbagens. Thus, at concentrations that induce endocycling, we confirmed that Aurora kinase inhibitors protect NSCLC cells from the cytotoxicity induced by all anti-mitotic cancer agents tested; but not agents targeting mechanisms functional in interphase. These endocycling NSCLC cells can re-enter the proliferative cell cycle if Aurora kinase inhibitors are withdrawn. We also demonstrate that chemical programming of the cells to endocycle with Aurora kinase inhibitors can serve as a methodology to identify antimitotic agents in a simple, high-throughput fashion.

## Results

### NSCLC cells exhibit three distinct dose response phenotypes to Aurora kinase A inhibitors

Aurora kinase A is an important regulator of mitosis and functions in the assembly of the mitotic spindle ^31–33^. Several different Aurora A inhibitors have been developed and are currently in clinical trials as anti-cancer drug candidates ^34^. In a previous study, we investigated these inhibitors and observed an unusual drug response relationship with an Aurora kinase A inhibitor, MLN8237/Alisertib (MLN8237 henceforth) ^35^. In contrast to the standard sigmoidal or monophasic dose-response phenotypes (Fig. 1a, c), a subgroup of cell lines treated with MLN8237 exhibited a multiphasic "camelback" drug-response pattern (Fig. 1e). VX-689, another Aurora kinase A inhibitor, also caused a similar response pattern in the same panel of non-small cell lung cancer (NSCLC) lines (Fig. 1b, d, f).

**Figure 1.**
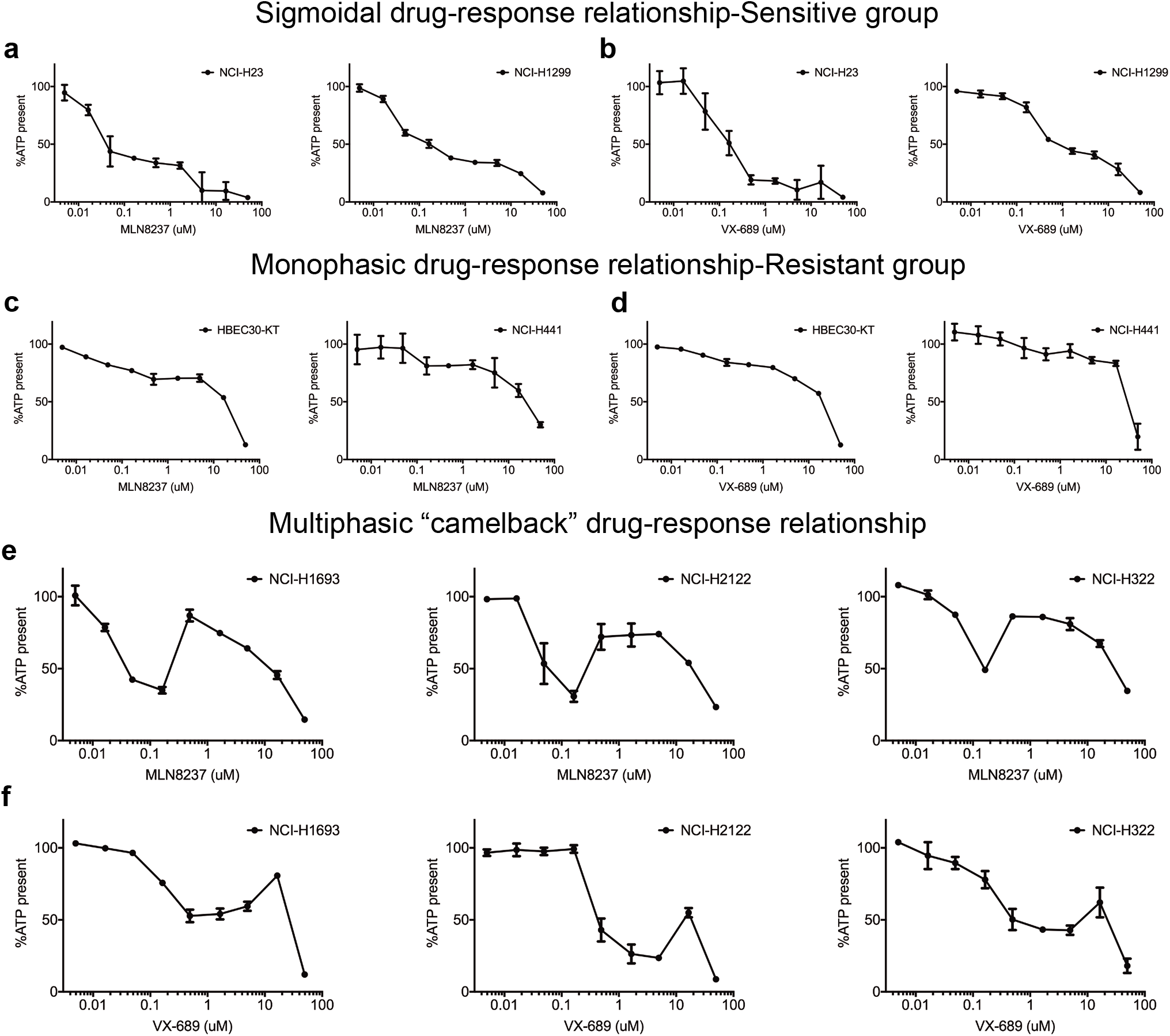
There are distinctly different response phenotypes to Aurora Kinase inhibition. **a-f**, Cells were treated with Aurora A selective inhibitors MLN8237 or VX-689 at the concentrations shown and cell viability was measured with a CellTiter-Glo assay, which measures cellular ATP as an indicator of living cells. Data are means of triplicate biological replicates and error bars are standard deviations (s.d). Some error bars are smaller than the data symbols. Cell lines representative of three distinct dose response curves are shown.

This unexpected drug response pattern of increased cell survival at higher drug concentrations suggested that secondary targets of these inhibitors might control a mechanism leading to drug resistance. This would be of concern, since peak plasma concentrations of MLN8237 in humans reach 3 μM, well above the concentration at which it selectively inhibits its primary target ^36^. Thus, it was of interest to identify the secondary targets of Aurora A inhibitors and the respective drug resistance mechanism.

### Inactivation of Aurora kinases in combination restores cell survival and growth

NSCLC line NCI-H1693 was selected as representative of cell lines capable of this unusual dose-response because it had the most dramatic phenotype with 35% of the cells growing at 87% of the rate of an untreated population in the presence of higher, potentially non-selective concentrations of MLN8237 (Fig. 2a). NCI-H1693 cells had a first phase EC50 of 23 nM for MLN8237 and 64 nM for VX-689 (Fig. 2a, c). The EC50s of MLN8237 and VX-689 for the second, "restorative" phase in which the viability increased, was 180 nM and 26 μM, respectively. At the highest and clinically irrelevant concentrations (Phase 3), MLN8237 gains a "general toxin" characteristic, which was not investigated in this study. In parallel, we measured Caspase 3/7 activity to determine if apoptotic cell death occurred in the presence of MLN8237 and VX-689 at any of concentrations measured. Apoptosis was first significantly detected at 16 nM for MLN8237 and 47 nM for VX-689. However, caspase activation was not detected during the second phase of the response to each drug (Fig. 2b, d). In an orthogonal assay for cell growth, NCI-H1693 cells were stained with crystal violet after seven days treatment with either MLN8237 or VX-689 and cell densities were compared with the density of cells stained on day 0. We found that growth of NCI-H1693 cells was inhibited at Aurora A-selective concentrations and was restored at 410 nM and 33.3 μM of MLN8237 and VX-689, respectively (Fig. 2e, g). Thus, the second phase of response to these two inhibitors not only helped NCI-H1693 cells survive, but also restored their growth.

**Figure 2.**
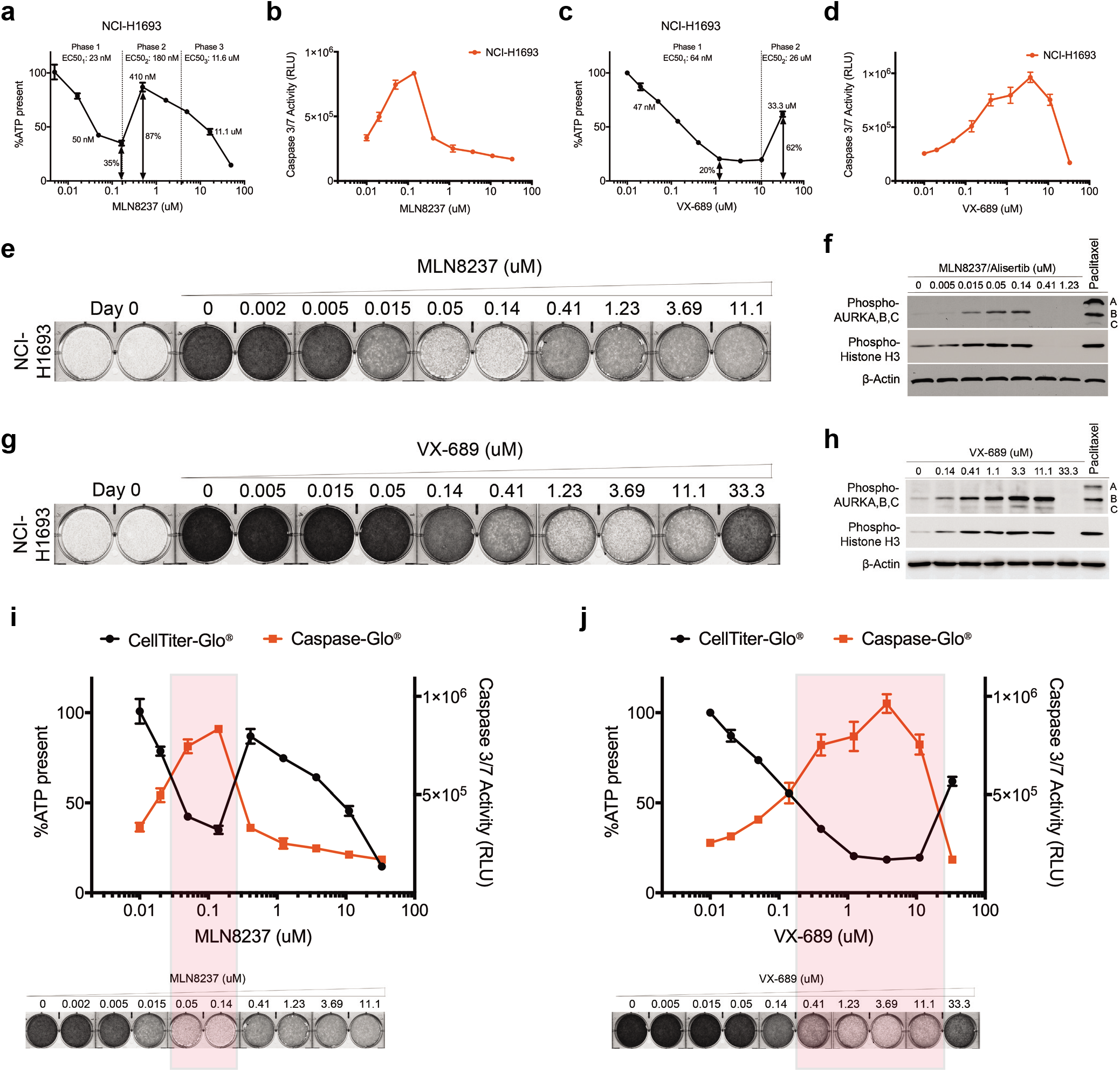
Aurora A inhibitors at higher concentrations antagonize their own cytotoxic activity and restore growth. **a,** NCI-H1693 cells were treated with serial concentrations of MLN8237 for four days and cell viability was measured with a CellTiter-Glo assay, which measures ATP. **b,** NCI-H1693 cell line was treated with serial concentrations of MLN8237 for two days and active apoptosis was measured with a CaspaseGlo 3/7 assay. **c, d,** CellTiter-Glo and CaspaseGlo 3/7 assays were repeated with VX-689 treatments. Data are means of triplicate biological replicates with s.d. Some error bars are smaller than the data symbols. **e,** NCI-H1693 cells were treated with serial concentrations of MLN8237 or **g,** VX-689 for seven days and cell growth quantified with a crystal violet staining assay. **f,** Cell lysates of NCI-H1693 cell line were collected 48-hours after treating with MLN8237 or **h,** VX-689 and immunoblotted to measure phosphorylated histone H3 and phosphorylated Aurora A, B, and C. Paclitaxel was used in both sets of immunoblots as positive control and B-Actin was used as internal immunoblotting control. **i, j,** Overlay of cell viability, caspase 3/7 activity and crystal violet staining images with immunoblotting results at Aurora A kinase selective and non-selective (pan-Aurora kinase) inhibitory concentrations of MLN8237 and VX-689.

Kinase inhibitors frequently are selective for their intended target only over a narrow concentration range ^37–39^. Aurora kinases B and C are two orthologues of Aurora A in humans and their high degree of homology is a well-known challenge for creating inhibitors specific to Aurora kinase A. Therefore, Aurora B and Aurora C were obvious candidates for the proteins responsible for reversing cytotoxic effects at higher concentrations of Aurora A inhibitors. Aurora kinases auto-phosphorylate when bound to their activators and their phosphorylation levels directly indicate their activation status in cells. In addition, Histone H3 is a well-characterized substrate of Aurora kinases B and C, and its phosphorylation at Serine 10 is a direct reporter of Aurora B or C activity ^40–43^. For these reasons, the phosphorylation statuses of Aurora kinases A, B, C and Histone H3 were measured 48 hours after treating NCI-H1693 cells with MLN8237 or VX-689 at a series of concentrations covering their two distinct phases of response. In these experiments, Paclitaxel at its EC100 (20 nM) was used as a positive control to inhibit the cell cycle at the mitotic stage where Aurora kinases are active. We observed increased phosphorylation of Aurora B and C, and Histone H3 starting at concentrations 50 nM of MLN8237 and 140 nM of VX-689. However, starting at 410 nM of MLN8237 and 33.3 μM of VX-689, phosphorylation of Aurora B and C, and Histone H3 was completely undetectable (Fig. 2f, h). Thus, there was a precise overlap between cell death, apoptosis and selective inhibition of Aurora A and between lack of apoptosis, continued cell growth and inhibition of Aurora B and C (Fig. 2i, j).

To test the hypothesis that Aurora B and C are the secondary targets causing the increase in survival and growth of cancer cells, we used two selective Aurora B and C inhibitors, AZD1152/Barasertib (AZD1152 henceforth) and GSK1070916 ^44,45^. We first verified their selectivity for Aurora B and C compared to Aurora A by measuring auto-phosphorylation levels of Aurora kinases A, B and C, and phosphorylation of Histone H3 in NCI-H1693 cells. Paclitaxel at its EC100 was used as a positive control. At 50 nM AZD1152 or 15 nM GSK1070916 auto-phosphorylation of Aurora B, C and Histone H3, was not detected; but Aurora A was phosphorylated (Fig. 3a, b). This indicated that at those concentrations, NCI-H1693 cells were arrested in mitosis at a point after activation of Aurora A. However, starting at 1.23 μM for AZD1152 or 140 nM for GSK1070916, phosphorylation of Aurora A was not detected, indicating that both drugs also inhibit Aurora A at higher concentrations.

**Figure 3.**
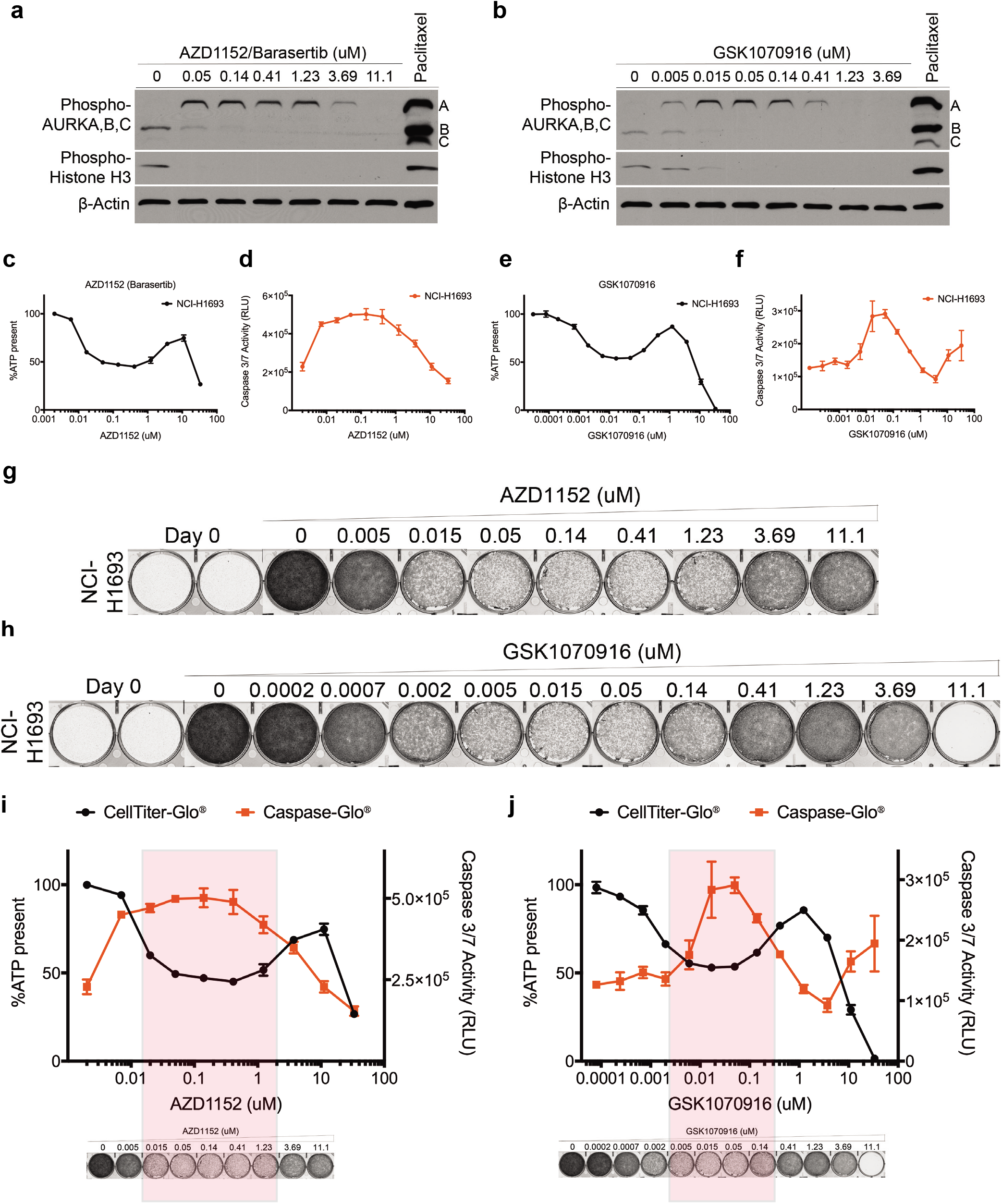
Aurora B, C inhibitors exhibit a camelback drug response pattern similar to Aurora A inhibitors. **a,** Cell lysates of NCI-H1693 cells were collected 48-hours after treating with AZD1152 (Barasertib) or **b,** GSK1070916 and immunoblotted to monitor phosphorylated histone H3 and phosphorylated Aurora A, B, C levels. Paclitaxel was used in both sets of immunoblots as positive control and B-Actin was used as internal immunoblotting control. **c,** NCI-H1693 cells were treated with serial concentrations of AZD1152 for four days and cell viability was measured with a CellTiter-Glo assay. **d,** NCI-H1693 cells were treated with serial concentrations of AZD1152 for two days and cell death was measured with a CaspaseGlo 3/7 assay. **e, f,** Assays were repeated with GSK1070916 treatments. Data are means of triplicate biological replicates with s.d. Some error bars are smaller than the data symbols. **g,** NCI-H1693 cell line was treated with serial concentrations of AZD1152 or **h,** GSK1070916 for seven days and cell growth was measured with a crystal violet staining assay. **i, j,** Overlay of cell viability, caspase 3/7 activity and crystal violet staining images with immunoblotting results at selective and non-selective concentrations of AZD1152 and GSK1070916.

The dose-response pattern of viability of NCI-H1693 cells to AZD1152 and GSK1070916 resembled the multiphasic "camelback" response of those cells to Aurora A selective inhibitors MLN8237 and VX-689 (Fig. 3c-f). AZD1152 and GSK1070916 each caused significant reduction in viability of NCI-H1693 cells at concentrations selective for Aurora B and C. However, growth was restored when NCI-H1693 cells were treated with concentrations above 1.23 μM and 140 nM of AZD1152 and GSK1070916, respectively (Fig. 3g-h), where they began to antagonize Aurora A as a secondary target. These data suggested that inhibiting Aurora A or Aurora B and C activities alone caused significant mitotic catastrophe leading to cytotoxicity; but inhibition of all three Aurora kinases together did not (Fig. 3i, j).

To ensure that Aurora kinases A, B and C, but not a previously unrecognized fourth target, are solely responsible of restoring viability induced with individual treatments of each of these four inhibitors, we treated NCI-H1693 cells with these four inhibitors as doublet combinations at concentrations where they are selective for their primary targets (Fig. 4a, b). Combination therapy of Aurora A and Aurora B, C inhibitors did not cause additive or synergistic cytotoxicity; but instead allowed cell survival and growth (Fig. 4c, e). We measured the activities of Caspase 3/7 (Fig. 4d, f) and used crystal violet staining (Fig. 4g, i) to assess apoptosis and cell growth. Inhibiting the Aurora kinases together allowed NCI-H1693 cells to survive and grow whereas inhibiting Aurora A or B selectively was inhibitory. Consequently, concentrations of Aurora B, C inhibitors selective for their target could prevent toxicity by Aurora A inhibitors when combined together in NCI-H1693 cells (Fig. 4h, j).

**Figure 4.**
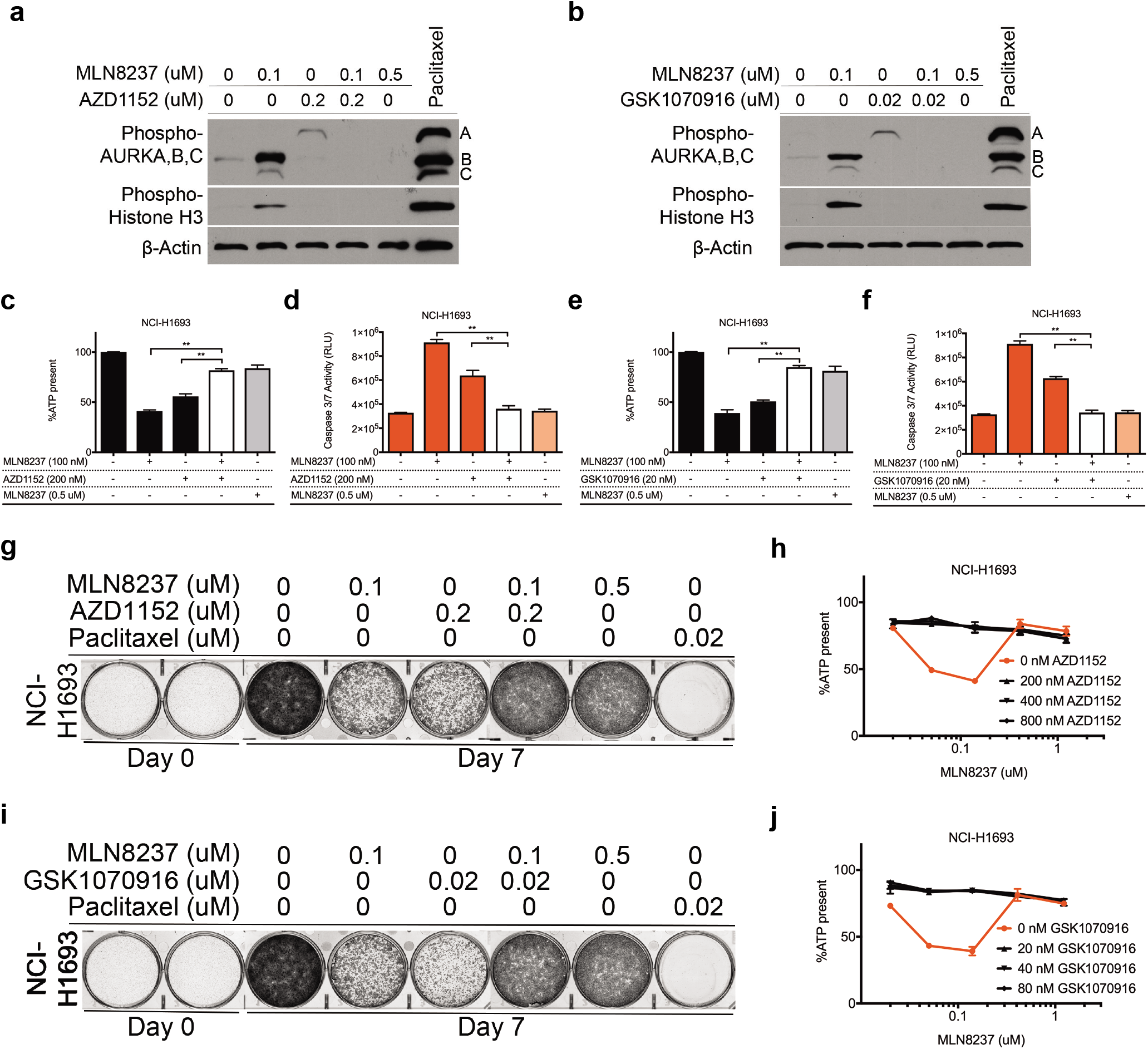
Combining Aurora A and B, C inhibitors at concentrations selective for their primary targets prevents cytotoxicity and allows cell growth. **a,** Cell lysates of NCI-H1693 cells were collected 48-hours after treating with combinations of MLN8237 and AZD1152 or **b,** MLN8237 and GSK1070916 and immunoblotted to detect phosphorylated histone H3 and phosphorylated Aurora A, B, C. Paclitaxel was used in both sets of immunoblots as a positive control and β-Actin was used as an internal immunoblotting control. **c,** NCI-H1693 cells were treated with combinations of MLN8237 and AZD1152 at concentrations selective for their primary targets for four days and cell viability was measured with a CellTiter-Glo assay. **d,** NCI-H1693 cells were treated with combinations of MLN8237 and AZD1152 at their selective for two days and cell death was measured with a CaspaseGlo 3/7 assay. **e, f,** Identical assays were repeated with GSK1070916 treatments. **g,** NCI-H1693 cells were treated with combinations of Aurora A and B/C selective inhibitors for seven days and cell growth was measured with a crystal violet staining assay. **h,** NCI-H1693 cells were treated with serial concentrations of MLN8237 in combination with serial dilutions of AZD1152 for four days and cell viability was measured with a CellTiter-Glo assay. **i, j,** Identical assays were repeated with GSK1070916 treatments. Data from CellTiter-Glo and CaspaseGlo 3/7 assays are presented as means of triplicate biological replicates with s.d. Some error bars are smaller than the data symbols.

Based on these observations, we hypothesized that pan-Aurora kinase inhibitors would be non- or minimally effective on NCI-H1693 cells when compared with the first response phases of MLN8237, VX-689, AZD1152 and GSK1070916. We performed biochemical assays identical to the ones performed with the four Aurora A or B and C selective inhibitors. We observed that pan-Aurora inhibitors, VX-680/Tozasertib (VX-680 henceforth) and PHA-739358/Danusertib (PHA-739358 henceforth) have a resistance response pattern resembling the second phase of multiphasic response of the first four selective inhibitors (Extended Data Fig. 1a-h).

To determine how frequently NSCLC cell lines survived simultaneous inhibition of all Aurora kinases, we measured the dose-responses to MLN8237 in a panel of 77 NSCLC and 4 immortalized human bronchial epithelial cell (HBEC) lines. More than fifty percent of these cell lines had a multiphasic camel-back drug response pattern with more cell survival and growth at the pan-Aurora kinase inhibitory phase in comparison to the responses at lower Aurora A-selective doses (Fig. 5a). In addition, the rest of tested NSCLC lines were either uniformly sensitive or uniformly resistant to the effective doses of MLN8237 with a sigmoidal or monophasic response, respectively (Fig. 5a). An identical screen with VX-689 produced results similar to those obtained with the screen of MLN8237 (Fig. 5b). In these studies, Erlotinib, an EGFR-targeted therapy agent and FDA-approved drug for NSCLCs, was used as control in parallel to MLN8237 and VX-689, on the same day at identical conditions and no multiphasic "camelback" response pattern was observed (Fig. 5c). Overall, our data indicated that the multiphasic response pattern against Aurora A inhibitors is common for NSCLC cell lines, and more cells survived and continued to grow when all Aurora kinases were inhibited in comparison to selective Aurora A inhibition.

**Figure 5.**
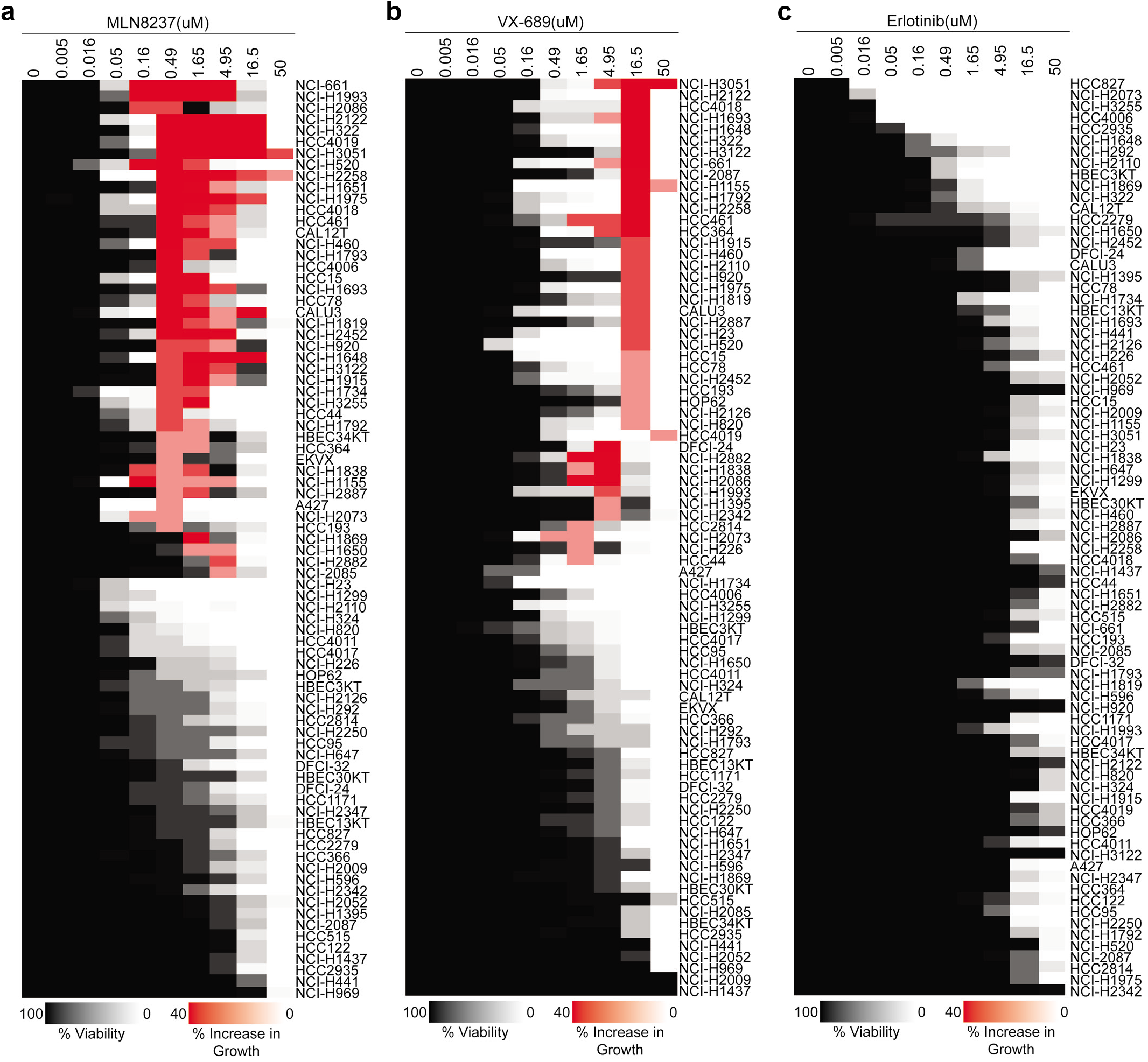
Survey of a NSCLC line panel reveals that more than 50% of cell lines have more cancer cell survival and growth at higher concentrations of Aurora kinase A inhibitors. **a,** A panel of NSCLC and HBEC lines was treated with MLN8237, **b,** VX-689 or **c,** Erlotinib. The results of dose–response experiments are shown in heatmap format. Decrease in viability in comparison to the untreated control is shown in shades of gray and white. Increase in viability in comparison to the cytotoxicity induced at lower concentrations is colored in shades of red.

### Inactivation of Aurora kinase signaling allows cells to develop into polyploid multinucleated (PP/MN) giant cancer cells

To determine what type of growth cells adopt after the loss of Aurora kinase signaling, we examined cellular morphology and DNA content after inhibiting Aurora kinases individually or in combination. After the inhibition of Aurora kinases with the non-selective high concentration of MLN8237, we observed that NCI-H1693 cells grow substantially in size instead of numbers and therefore, cellular ATP levels increase similar to untreated proliferating cells (Fig. 6a-b). Their nuclear morphology together with enlarged size and elevated DNA content, demonstrated that these cells develop into polyploid multinucleated (PP/MN) giant cancer cells (Fig. 6b), a cancer cell type frequently found in drug-resistant tumor populations and high-grade tumors. The average area of cells treated with pan-Aurora kinase inhibitor for four days increased and cells had much more nuclear DNA in comparison to the untreated ones (Fig. 6c). Polyploid genome was also compartmentalized in multiple nuclei in these cells (Fig. 6c). Cell numbers were not increasing (Fig. 6d) in the treated samples, but were balanced by increasing the cellular size. Therefore, total ATP levels present in the treated wells remained similar to the wells with untreated cells. DNA replication also continued (Fig. 6d). Increasing DNA content and cell size are characteristics of an endocycle ^46^. Overall, our findings suggested that after inhibiting Aurora kinase signaling, cells might omit all or most of mitosis, continue with intact Gap (G) and Synthesis (S) phases of cell cycle, and develop into PP/MN giant cancer cells (Fig. 6e).

**Figure 6.**
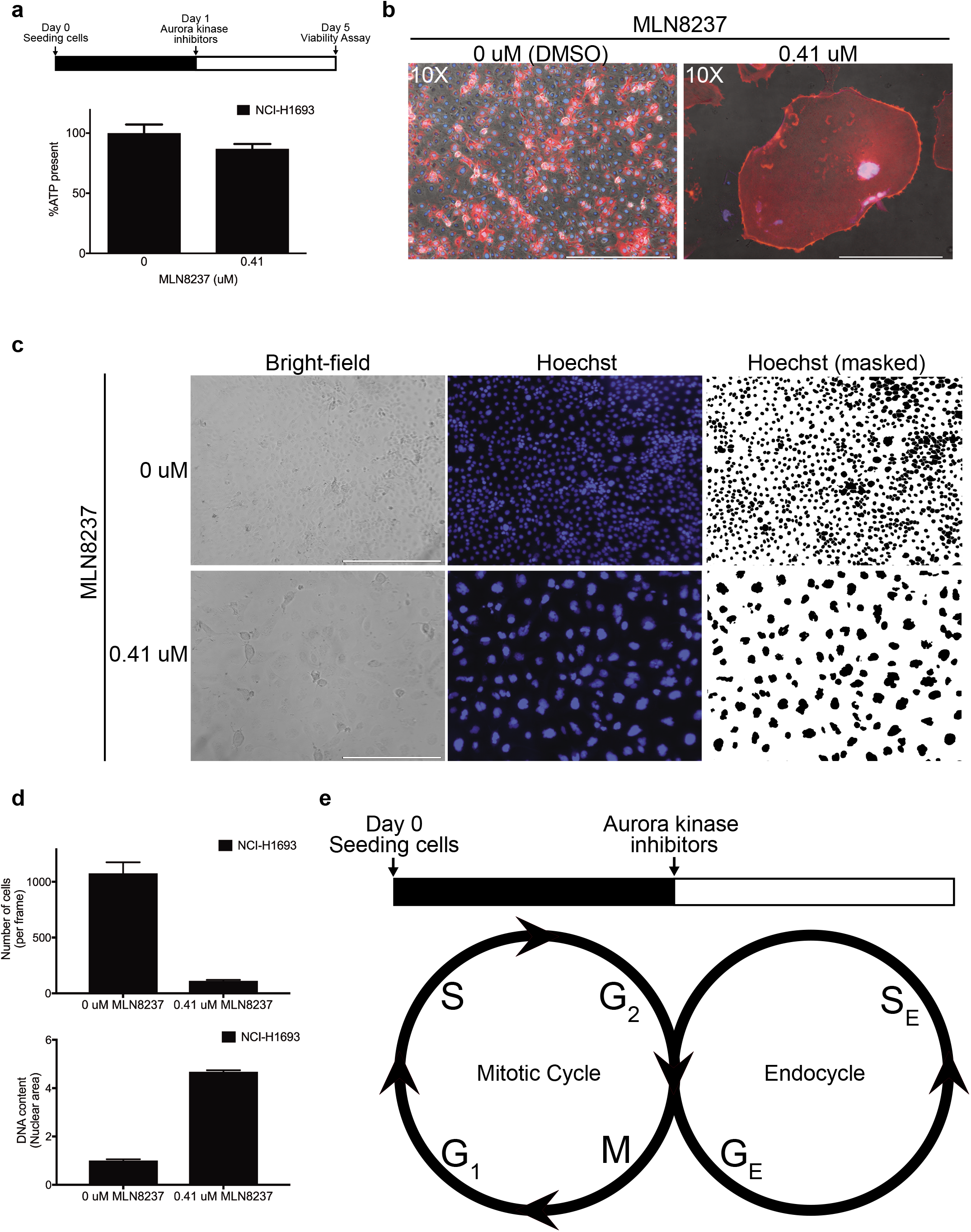
Loss of Aurora kinase signaling allows cells to grow into polyploid multinucleated giant cancer cells. **a,** NCI-H1693 cells were treated with 0.41 μM of MLN8237 for four days and the ATP present was measured with a CellTiter-Glo assay. **b,** NCI-H1693 cells were treated with DMSO or 0.41 μM of MLN8237 for 20 days, stained with Phalloidin (red) and Hoechst (blue), and visualized at 10X magnification (Scale bar, 400 μM). **c,** NCI-H1693 cells were treated with 0.41 μM of MLN8237 for four days and visualized with bright-field or fluorescent microscopy at 10X magnification. Hoechst reagent was used to visualize the nuclei (Scale bar, 400 μM). **d,** The numbers of cells and their mean nuclear area from treated and untreated control samples are graphed. **e,** A proposed model of an endocycle induced with the loss of Aurora kinase signaling.

### Induced PP/MN giant cancer cells resist anti-mitotic agents

If an endocycle variant avoids mitosis when Aurora kinase signaling is turned off, we hypothesized that mitotic machinery including spindle-related components and regulators should partially or completely be dispensable in the induced PP/MN giant cells. Thus, cells would be non-responsive to anti-mitotic agents under these conditions instead of causing additive or synergistic effects whereas perturbing interphase cellular machineries should remain cytotoxic (Fig. 7a). We tested NCI-H1693 cells with 17 anti-cancer or cytotoxic agents including 9 acting during interphase (Gemcitabine, Carboplatin, Cisplatin, Etoposide, Topotecan, Doxorubucin, Pemetrexed, Methotrexate, 5-Fluorouracil, Aphidicollin and Bafilomycin A) and 8 inhibiting mitosis (Paclitaxel, Docetaxel, Vinorelbine, Vinblastine, PLK1 inhibitor BI-6727 (Volasertib), CENPE inhibitor GSK923295, EG5 inhibitors Ispinesib and VS-83). For the cell viability assessments, the EC100 of each agent was used (Extended Data Table 1). Aurora kinase inhibition prevented toxicity from all anti-mitotic drugs whereas cells were sensitive to interphase perturbagens regardless of the presence of Aurora inhibitors (Fig. 7b). In addition to individual treatments with these anti-cancer agents, we also added a cocktail of the eight anti-mitotic agents to our survey and observed that PP/MN giant cancer cells generated by pan-Aurora kinase inhibition were able to avoid death even in that extreme condition (Fig. 7b). In parallel to the cell viability assays, we stained NCI-H1693 cells with crystal violet to check the status of cell growth on day 7 in comparison to the cell density on Day 0. Our results confirmed that NCI-H1693 cells survived and continued to grow in the presence of all anti-mitotics and their cocktail, but not the anti-interphase perturbagens (Fig. 7c). Lastly, we quantified the extent of tolerance of the PP/MN giant cells to individual anti-mitotic agents by performing cell viability assays with serial concentrations of eight anti-mitotic drugs in the presence and absence of pan-Aurora kinase inhibitors. We observed at least 4 logarithmic (log) magnitude of drug resistance to Paclitaxel, 5 logs to Docetaxel (Fig. 7d), and at least 2 logs of significant resistance to others (Extended Data Fig. 2a-f).

**Figure 7.**
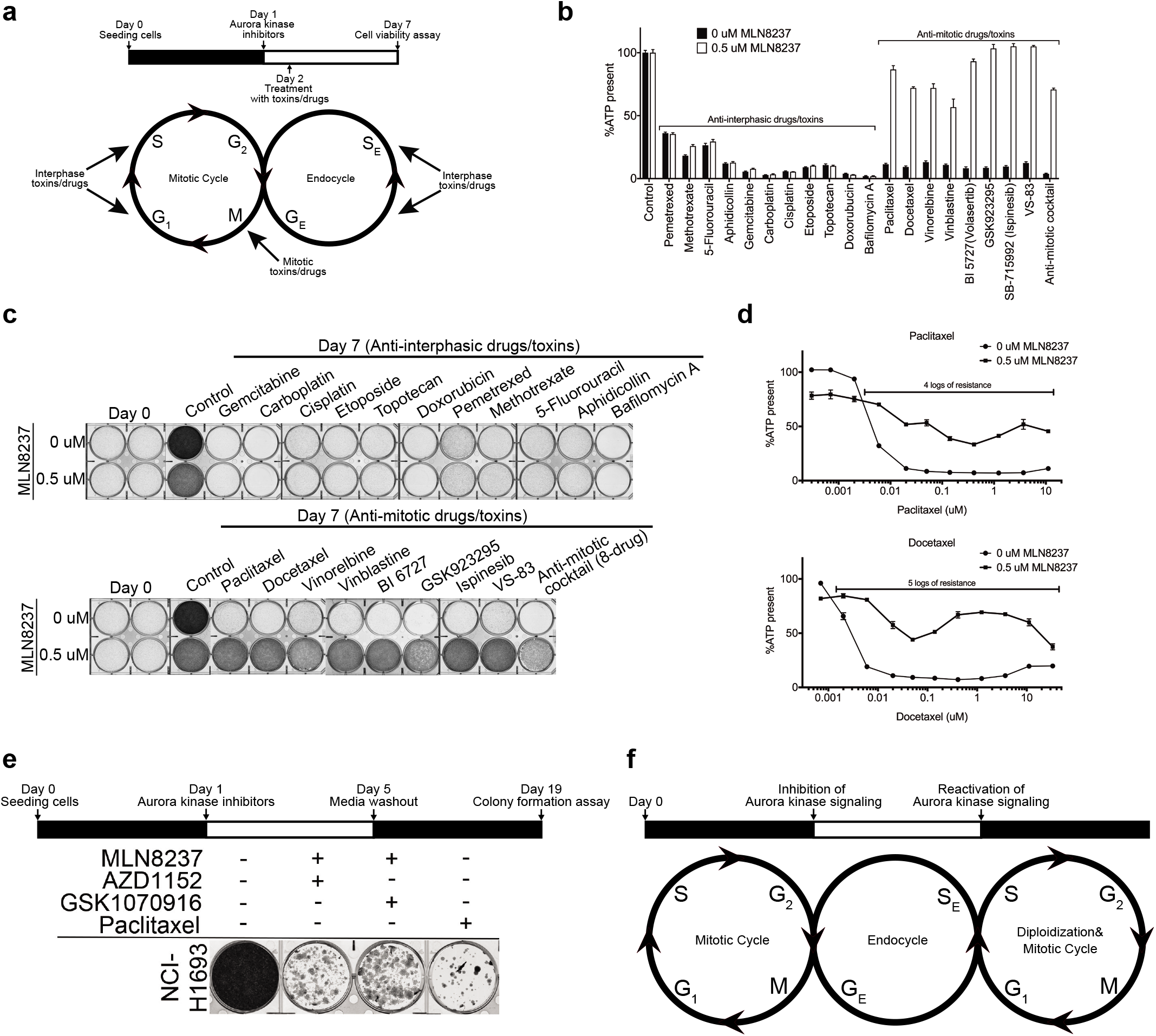
Inhibition of Aurora kinase signaling provides resistance to anti-mitotic toxins, but not interphase toxins. **a,** NCI-H1693 cells were co-treated with DMSO or 0.5 μM of MLN8237 and IC100 concentrations of interphase or mitotic perturbagens. **b,** Cell viability and growth were assessed with a CellTiter-Glo assay (after four days of treatment) or **c,** crystal violet staining assay (after seven days of treatment), respectively. **d,** NCI-H1693 cells were co-treated with DMSO or 0.5 μM of MLN8237 and serial concentrations of Paclitaxel or Docetaxel. Cell viability was measured with a CellTiter-Glo assay. **e,** NCI-H1693 cell line was treated with 0.1 μM of MLN8237 and 0.2 μM AZD1152 or 20 nM GSK1070916 for four days. Treatment media were replaced with drug-free media for 14 days. Colony formation was measured with a crystal violet staining assay. Data of CellTiter-Glo assay are means of triplicate biological replicates with s.d. Some error bars are smaller than the data symbols. **f,** Proposed model of the cellular life cycle in the presence and absence of Aurora kinase signaling.

Finally, we investigated whether the NCI-H1693 PP/MN cells that had switched to endocycle could reenter the proliferative cell cycle and regenerate daughter cells. Four days after inhibiting Aurora kinase signaling with combinations of Aurora A inhibitor, MLN8237 and Aurora B, C inhibitors AZD1152 or GSK1070916 at concentrations selective for their primary targets, the media with the combination treatments was washed out and NCI-H1693 cells were allowed to grow in the absence of any drug treatments for 14 days. NCI-H1693 cells recovered from the combination treatment and were able to form colonies, indicating the ability of cells to reenter mitotic cell cycle (Fig. 7e). In comparison, NCI-H1693 cells treated with Paclitaxel had significantly fewer colonies. These observations indicate that loss of Aurora kinase signaling initiates an endocycle, which maintains growth in cell size instead of increases in cell numbers. These cells grow into PP/MN giant cancer cells and can re-enter a proliferative cell cycle when Aurora kinase signaling is restored (Fig. 7f).

Identification of the target and mechanism of action for biologically active molecules remains a major task for drug discovery. To explore the possible use of a chemically-induced endocycle as a high-throughput assay to identify antimitotic chemicals, NCI-H2122 cells, one of the NSCLC lines with the most dramatic multiphasic camelback response to MLN8237 (Fig. 1e), was used to profile 12 cytotoxic chemicals in the presence or absence of 0.5 uM of MLN8237, a concentration that induces endocyling (Fig. 8a, b). The chemicals used in this test had been identified as toxic for subsets of our NSCLC cell line panel in a previous high-throughput screen, but their mode of action was unknown ^47^. Of the 12 chemicals, MLN8237-induced NCI-H2122 PP/MN giant cells were protected from one, SW172170. To determine if SW172170, but not the other chemical probes, had an anti-mitotic mechanism of action as predicted by our assay, 48 hours after treating with the compounds, we checked for increased phosphorylation of Histone H3 as a sign of mitotic arrest. SW172170 was the only compound showed increase in Histone H3 phosphorylation levels (Fig. 8d). As an orthogonal assay, NCI-H2122 cells treated with SW172170 as a single agent were stained for DNA and tubulin. These cells had condensed DNA and multipolar spindles, which are the typical characteristics of mitotic arrest and catastrophe (Fig. 8e). This demonstrates that chemical inhibition of Aurora kinases in a cell line that responds by endocyling can be used as a screen for anti-mitotic mode of action and therefore, can be employed as a tool to assess the cell cycle phase (interphase vs mitosis), in which a toxic chemical acts.

**Figure 8.**
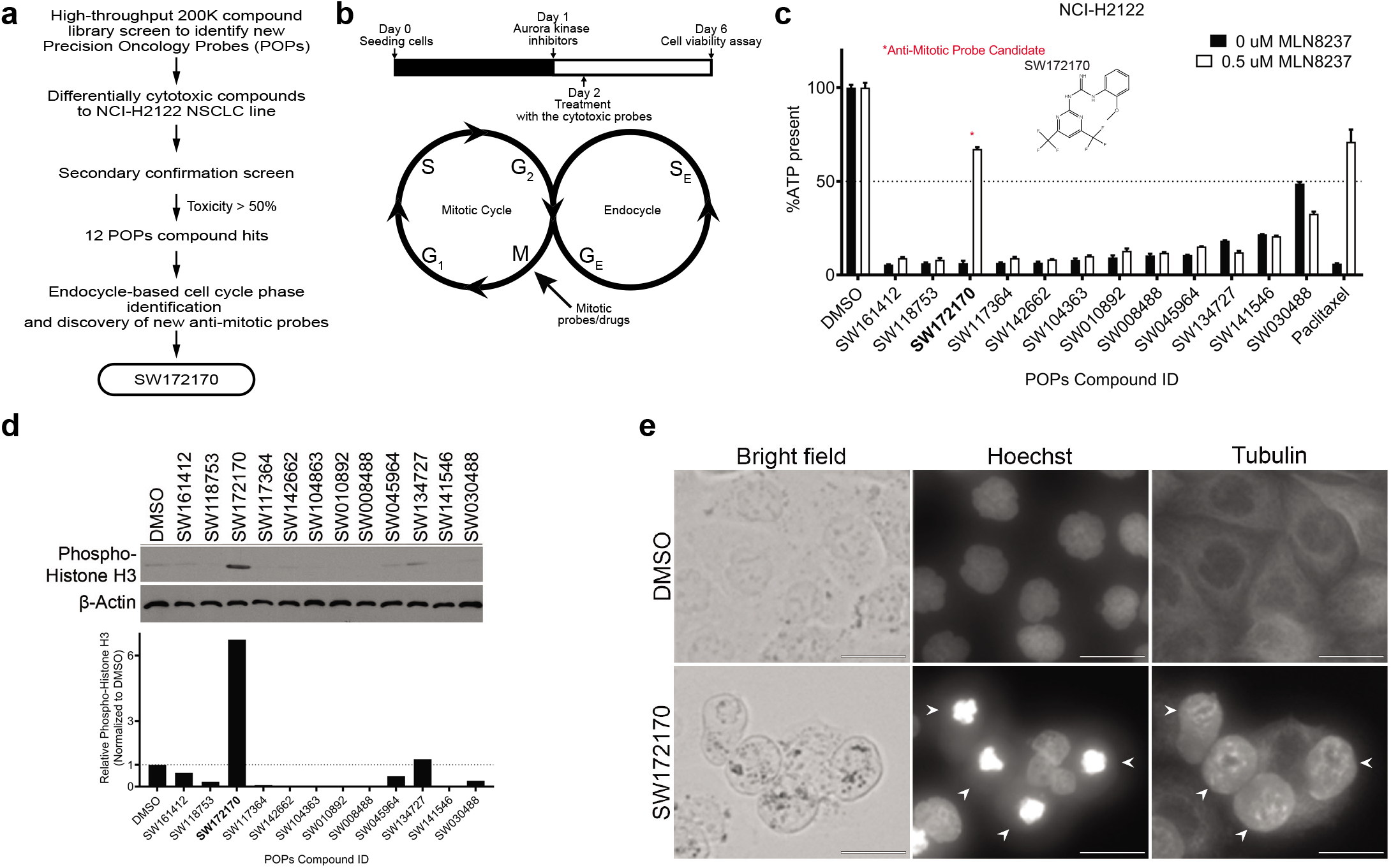
Induction of polyploid multinucleated giant cancer cells through chemical inhibition of Aurora kinase signaling can serve as a strategy to identify anti-mitotic chemicals. **a,** Flow chart of the drug discovery project designed to identify probes selectively cytotoxic to NCI-H2122 cells. **b,** Proposed model for the anticipated outcome of screen identifying new anti-mitotic perturbagens. **c,** NCI-H2122 cells were treated with DMSO or 0.5 μM of MLN8237 and each of 12 chemical hits identified in a compound library screen. Cell viability was measured with a CellTiter-Glo assay. Paclitaxel was used as positive control. Structure of SW172170, the anti-mitotic candidate probe is shown. **d,** Cell lysates of NCI-H2122 cell line were collected 48-hours after treating with each chemical and immunoblotted to measure phosphorylated histone H3. ß-Actin was used as internal immunoblotting control. Normalized band intensities of phosphorylated Histone H3 from treated and untreated control samples are graphed. **e,** NCI-H2122 cells were treated with SW172170 for four days and visualized with bright-field or fluorescent microscopy. Hoechst reagent and monoclonal antibodies against Tubulin were used to visualize the nuclei and mitotic spindles, respectively (Scale bar, 25 μM).

## Discussion

Tumors and tumor-derived cell lines contain polyploid giant cells with significantly increased genomic content, often within multiple nuclei. Their frequency increases after exposure to therapeutic agents. These PP/MN cells are not dormant, and form macroscopic colonies and tumors in immunodeficient mice ^8–10^. Their daughter cells harbor stemness, self-renewal and mitotic capacities ^11–13^. Consistent with these attributes, a significant number of recent reports with different biological systems (tumor-derived cell lines, animal models and specimens from cancer patients) have underlined the key roles played by PP/MN giant cancer cells in tumorigenesis, metastasis, immunosuppression and disease relapse after conventional cancer treatments ^3,4,48^. PP/MN cells contribute to solid tumor heterogeneity and are the main histological feature of malignant tumor in pathologic diagnosis. Our findings demonstrate that inactivation of Aurora kinases A, B and C combinatorially is a signal to switch to an endocycle that forms PP/MN giant cancer cells.

Our results indicate that in NSCLC cells Aurora kinases function together to keep the cancer cell in a proliferative cell cycle. In many cell types, only Aurora A and B kinases are expressed and the extent to which Aurora C plays a role in this activity is not determined by our work. In a majority of NSCLC cell lines we tested, chemical inhibition of all family members of Aurora kinases allowed the cells to adopt an alternative cell cycle where cells continue to replicate DNA, become multinucleate and grow in size, but do not divide. In this state, cells are resistant to all anti-mitotic drugs tested because the mitotic machinery is no longer engaged. Our findings together suggest that Aurora kinases may operate as a cell cycle switch and their off status generates PP/MN giant cancer cells tolerant to anti-mitotic therapy in those cell lines where the alternative cell cycle is available.

In nature there are numerous examples of alternative cell cycles, called endocycles (also known as endoreplication, endoreduplication or endomitosis), that generate healthy polyploid cells for specialized functions. Endocycling is essential for the proper development of specialized cell types in mammalian systems like megakaryocytes, placental trophoblast cells, hepatocytes and cardiomyocytes ^49^. The level of ploidy varies dramatically among tissues, organs and species. Mammalian placental trophoblast cells can exhibit ploidy levels of 4096 times the haploid number of chromosomes (4096C) ^50,51^. Megakaryocytes generate nuclei with DNA content up to 128C ^52^. The highest recorded level for hepatocytes and cardiomyocytes is 32C. By adulthood and with ageing, up to 70% of human and 85% of rodent cardiac myocytes are polyploid ^53,54^. The highest polyploidy level recorded in any animal is ~500,000C, reported from the silk-producing glands of silk moth *Bombyx mori* ^55^. The neurons of mollusks exhibit remarkable variations of ploidy, ranging from a low 32C in the land snail *Triodopsis divesta* to an astonishing 200,000C in the giant neurons of the sea hare *Aplysia californica* ^56,57^. A range of 4C to 64C is typical in plants although the highest level of polyploidy observed in plants is 24,576C in the endosperm of *Arum maculatum* ^58,59^. Cancers can take advantage of endocycles by creating heterogeneity of cell types, some of which are more drug resistant than the parental cells ^60^.

It is possible that some cancer cells do this by adopting an alternate cell cycle program similar to those available to specialized cell types. In the brain, a blood-brain barrier is formed by endothelial cells and these specialized cell types are known to be decorated with efflux pumps and upregulate drug metabolizing enzymes ^61,62^. The glial cells forming the barrier in fruit fly *Drosophila melanogaster* was shown to require polyploidization for proper differentiation ^63,64^. In cancers, drug resistance mechanisms to lower the cytosolic drug concentrations with these two systems are well-documented and could originate from cancer cells that were capable of growing in endothelial cell-like endocycles.

How mitotic cycles are molecularly routed towards endocycles is largely unknown. However, a recent study on cells from *Drosophila melanogaster* uncovered a role for Aurora kinase B in endoreplication and its morphogenesis^65^. Reduced expression of Aurora kinase orthologues in plants was also shown to control the switch from meristematic cell proliferation to endoreduplication and differentiation^66^. Furthermore, Aurora kinase A was found downregulated during the megakaryocyte endomitosis and differentiation in mice^67^. In our experiments, chemically inhibiting Aurora kinases A or B alone did not induce endocycling in human lung cancer cells, but led to growth arrest and reduced cell numbers. Whether this is a difference between animal, plant and insect cells, or cancer and normal cells remains to be determined. Unexpected adverse interactions in clinical trials with combination therapies occur and mostly the reasons remain unknown. Currently, 60 clinical trials with MLN8237, 10 with VX-689, 12 with AZD1152, 14 with GSK1070916, 11 with VX-680 and 8 with PHA-739358 have been reported and among those many are performed in combination with other anti-mitotic drugs such as Taxanes and Vinca alkaloids. Among the concluded trials, some revealed that targeted mitotic inhibitors, including Aurora kinase inhibitors, were less effective than expected ^68,69^. Secondly, enlargement of cells in non-responsive cancers or tumor models after treatments with various chemo- or -targeted therapies has been reported ^60^. Increased cell size was also observed in studies with Aurora kinase-targeted therapies including MLN8237 ^70–72^. However, the cause for this unexpected cell enlargement remained unknown. Our study presents a potential mechanistic explanation for these clinical results.

Lastly, the multiphasic dose-response pattern that we observed with inhibitors of Aurora kinase signaling is not appropriately recognized by the standard approaches for drug dose-response curve fitting. It was recently reported that the failure rate of the classical Hill equation is more than 25% in large scale screens ^73^. In experiments investigating the response of NSCLC cell panels to synthetic chemicals ^47^, we have observed that dose-responses poorly represented by sigmoidal curves are quite common. Some of these are undoubtably due to chemical solubility or other experimental artifacts, but our current report indicates that some indicate a biological response that is not considered by the assumptions underlying most commonly used curve fitting programs.

In summary, our data suggest that Aurora kinase signaling functions at the intersection between proliferative cell cycle and endocycle. Switching to endocycle variants creates growing polyploid multinucleated cancer cells tolerant to anti-mitotic therapy. PP/MN cancer cells also retain the capability to reenter the proliferative cell cycle after the treatments are terminated.

## Methods

### Cell lines

NSCLC and immortalized HBEC lines were generated by the laboratories of John Minna and Adi Gazdar and their identities have been confirmed by short tandem repeat analysis of cellular DNA (Promega Corp, Madison, WI, USA). All cell lines were determined to be mycoplasma free by testing with the e-Myco Plus Mycoplasma PCR Detection Kit (Bulldog Bio, 2523448). NSCLC lines were cultured in RPMI-1640 medium (GIBCO, 11875) supplemented with 10% fetal bovine serum (v/v) (Atlanta Biological, S11550). HBEC lines were grown in Keratinocyte-SFM medium (GIBCO, 17005) supplemented with EGF and keratinocyte extract (provided by the manufacturer). HBEC3-KT, HBEC30-KT and HBEC34-KT are HBECs immortalized with stable expression of CDK4 and hTERT ^74^.

### Reagents

MLN8237 (Alisertib) (ChemieTek, CT-M8237), VX-689 (Selleckchem, S2770), AZD1152 (Barasertib) (Selleckchem, S1147), GSK1070916 (Selleckchem, S2740), VX-680 (Tozasertib) (Chemietek, CT-VX680), PHA-739358 (Danusertib) (Selleckchem, S1107), Paclitaxel (LC Labs, P-9600), Docetaxel (Sigma-Aldrich, 01885), Vinorelbine detartrate salt hydrate (Sigma-Aldrich, V2264), Vinblastine sulfate (Sigma-Aldrich, V1377), BI 6727 (Volasertib) (Selleckchem, S2235), GSK923295 (Selleckchem, S7090), Ispinesib (Cayman Chemical Company, 18014), VS-83 (Calbiochem, 178273), Gemcitabine (ChemieTek, CT-GEM), Carboplatin (Sigma-Aldrich, C2538), Cisplatin (Sigma-Aldrich, 232120), Etoposide (LC Labs, E-4488), Topotecan (Tocris, 4562), Doxorubucin hydrochloride (Sigma-Aldrich, D1515), Pemetrexed (Tocris, 6185), Methotrexate (Tocris, 1230), 5-Fluorouracil (Sigma-Aldrich, F6627), Aphidicollin (Calbiochem, 178273), Bafilomycin A (Sigma-Aldrich, B1793), Erlotinib (LC Labs, E-4497) was purchased in powder form and stored at 10 mM or 20 mM stock concentrations in dimethyl sulfoxide (DMSO) at −20 °C. Phospho-AURKA (Thr288)/AURKB (Thr232)/AURKC (Thr198) (Cell Signaling, rabbit mAb, 2914, 1:1,000 dilution), Phospho-Histone H3 (Ser10) (Cell Signaling, rabbit mAb, 33770, 1:1,000 dilution) and β-Actin (MP Biomedicals, mouse mAb, 69100, 1:10,000 dilution) antibodies were used for immunoblotting assays. Texas Red™-X Phalloidin (ThermoFisher Scientific, T7471, 1:100 dilution), Tubulin antibody (Abcam, ab6160, 1:1000 dilution) and Hoechst 33342 reagent (ThermoFisher Scientific, H3570, 1:2000 dilution) were used for immunofluorescence assays.

### Microplate drug sensitivity assays

NCI-H1693 cells (3,000 per well) were seeded in 96-well plates and incubated at 37 °C for 24 h. Treatment with vehicle (DMSO) or serial dilutions of the indicated compounds in 9 concentrations from 1 to 33,300 nM were performed in triplicate (final DMSO concentration=0.34% in all wells). After 96 h, media were removed and cell viability was measured with a CellTiter-Glo assay (Promega, G7573).

A dose–response study of a panel of NSCLC and HBEC cell lines with MLN8237 (Alisertib), VX-689 and Erlotinib was performed in 384-well plate format. Cells were dispensed at a density of 1,000–1,500 cells per well with an automated liquid dispenser (BioTek MicroFlo; Thermo-Fisher, Inc.). Each concentration was screened in triplicate. Cells were incubated for 24 h at 37 °C in the presence of 5% CO_2_.with an automated liquid dispenser (BioTek MicroFlo; Thermo-Fisher, Inc.) Drugs/chemicals were dosed in DMSO at 12 concentrations ranging from 50 pM to 50 μM (half-log dilutions) in triplicate, using an Echo 555 acoustical transfer instrument. After incubating the cells at 37 °C for 96 h, cell viability was measured with a CellTiter-Glo assay. The comparative tests of three agents were performed on the same days, in identical conditions. Dose responses of each cell line to each agent were assessed at the indicated concentrations individually. Concentrations that caused increase in the measured viability after the initial cytotoxic effect of respective agent were colored in red.

### Microplate apoptosis assay

Treatments of the indicated compounds were performed in the same manner with the microplate drug sensitivity assays in 96-well plate format. 48 hours after treatments, culture media were removed by centrifuging the assay plates upside down in liquid-collecting containers at *30g* for 1 min. Caspase 3 and/or 7 activity was assessed with Caspase-Glo 3/7 assay purchased from Promega (G8090).

### Crystal violet staining assay

NCI-H1693 cells (1 × 10^5^ per well) were seeded in 6-well plates and incubated at 37 °C for 24 h. Treatment with vehicle (DMSO) or serial dilutions of the indicated compounds in 10 indicated concentrations were performed (final DMSO concentration=0.34% in all wells). After seven days, media were removed and cell growth was assessed with a crystal violet staining assay solution (6% glutaraldehyde, 0.5% crystal violet (w/v) in water).

### Western blotting

In all, 2 × 10^5^ cells were treated with the indicated compounds at indicated concentrations in six-well format. Forty-eight hours after treatments, the culture medium was aspirated and cells were washed with PBS. Cells were lysed in RIPA lysis buffer (50 mM Tris buffer pH 8, 150 mM NaCl, 1% NP-40, 0.5% sodium, 0.1% SDS, 0.5% Sodium deoxycholate) supplemented with protease and phosphatase inhibitors. The protein concentrations of cell lysates were measured with the Protein DC assay (Bio-Rad, 500-0111). Phosphorylation of histone H3 and Aurora kinases A, B, C, and β-Actin expression as internal control were monitored by immunoblotting with monoclonal antibodies.

### Light microscope and immunofluorescence assay

Cells in transparent bottom plates were fixed with 4% formaldehyde in PBS for 15 min at room temperature. After washing with PBS, cells were permeabilized with 0.25% Triton-X in PBS for 15 min at room temperature. After washing with PBS, cells were incubated with 10% Bovine Serum Albumin (BSA) blocking solution for 1 h. Next, cells were immunostained with Texas Red™-X Phalloidin or monoclonal Tubulin antibodies overnight. Hoechst 33342 reagent was used to stain DNA. Microscopic observations were performed on an EVOS FL Cell Imaging System. Mean nuclear area of cells after the treatments with the indicated compounds were calculated with the ImageJ software and graphed.

## Data Availability

The data supporting this study are available from the corresponding author upon reasonable request.

## Acknowledgements

We thank Bruce Posner and the personnel of the Simmons Comprehensive Cancer Center High-throughput Screening Core Facility for aid in dose-response experiments on the NSCLC cell line panel. The human tumor-derived cell panel was created by Drs. John D. Minna and Adi F. Gazdar. We thank our colleague Iryna Zubovych for critical comments. This work was supported by grants CA176284 and CA221978 to MGR.

## Author Information

### Affiliations

Departments of Biochemistry, UT Southwestern Medical Center, Dallas, TX, USA Vural Tagal, Michael G. Roth

Harold Simmons Comprehensive Cancer Center, UT Southwestern Medical Center, Dallas, TX, USA

Michael G. Roth

## Author Contributions

V.T. designed and performed the experiments. V.T. and M.G.R. conceived the study, analyzed data and wrote the manuscript.

## Extended Data Figure Titles and Legends

**Extended Data Figure 1.**
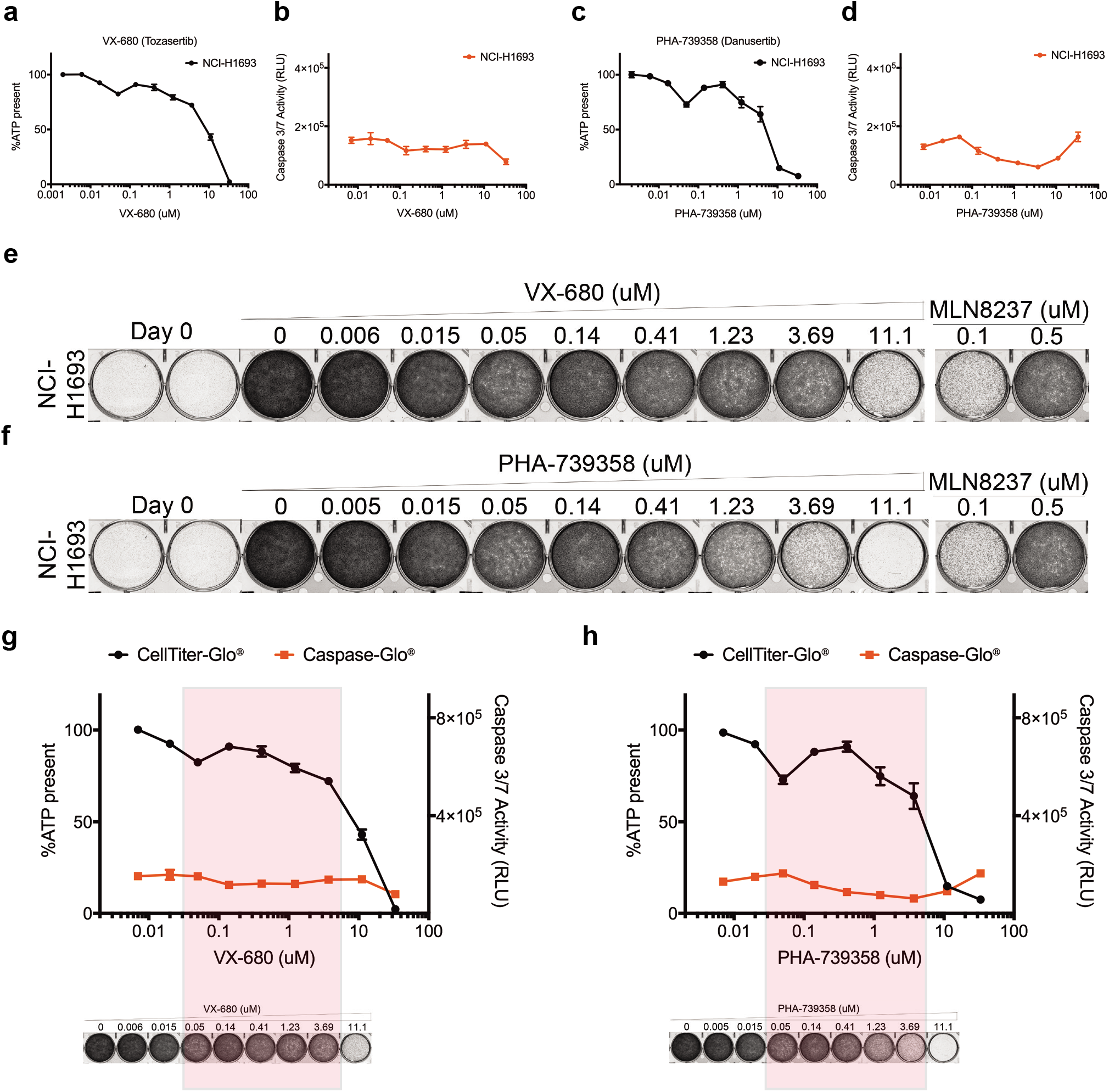
Pan-Aurora kinase inhibitors do not inhibit growth of NCI-H1693 cells. **a,** NCI-H1693 cells were treated with serial concentrations of VX-680 for four days and cell viability was measured with a CellTiter-Glo assay. **b,** NCI-H1693 cells were treated with serial concentrations of VX-680 for two days and apoptosis was measured with a CaspaseGlo 3/7 assay. **c, d,** Assays were repeated with PHA-739358 treatments. Data are means of triplicate biological replicates with s.d. Some error bars are smaller than the data symbols. **e,** NCI-H1693 cell line was treated with serial concentrations of VX-680 or **f,** PHA-739358 for seven days and cell growth was measured with a crystal violet staining assay. A concentration of MLN8237 specific for Aurora kinase A and an EC100 concentration of Paclitaxel were used as controls. **g, h,** Overlay of cell viability, caspase 3/7 activity and crystal violet staining images with immunoblotting results.

**Extended Data Figure 2.**
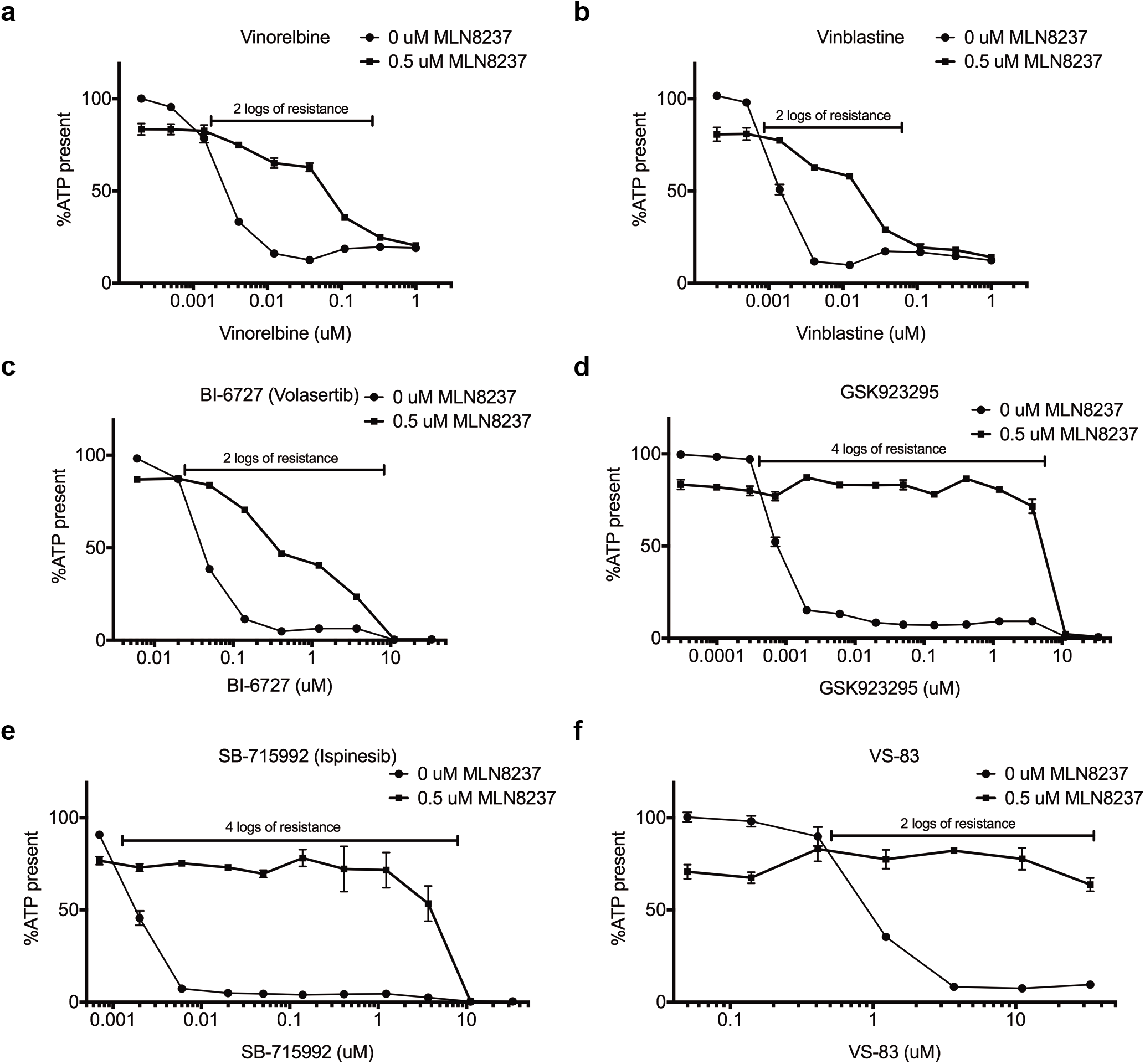
Endocycling cells resist anti-mitotic perturbagens. **a,** Vinorelbine, **b,** Vinblastine, **c,** BI 6727 (Volasertib), **d,** GSK923295, **e,** SB-715992 (Ispinesib) or **f,** VS-83 for four days and cell viability was measured with a CellTiter-Glo assay. Data are means of triplicate biological replicates with s.d. Some error bars are smaller than the data symbols.

**Extended Data Table 1.**
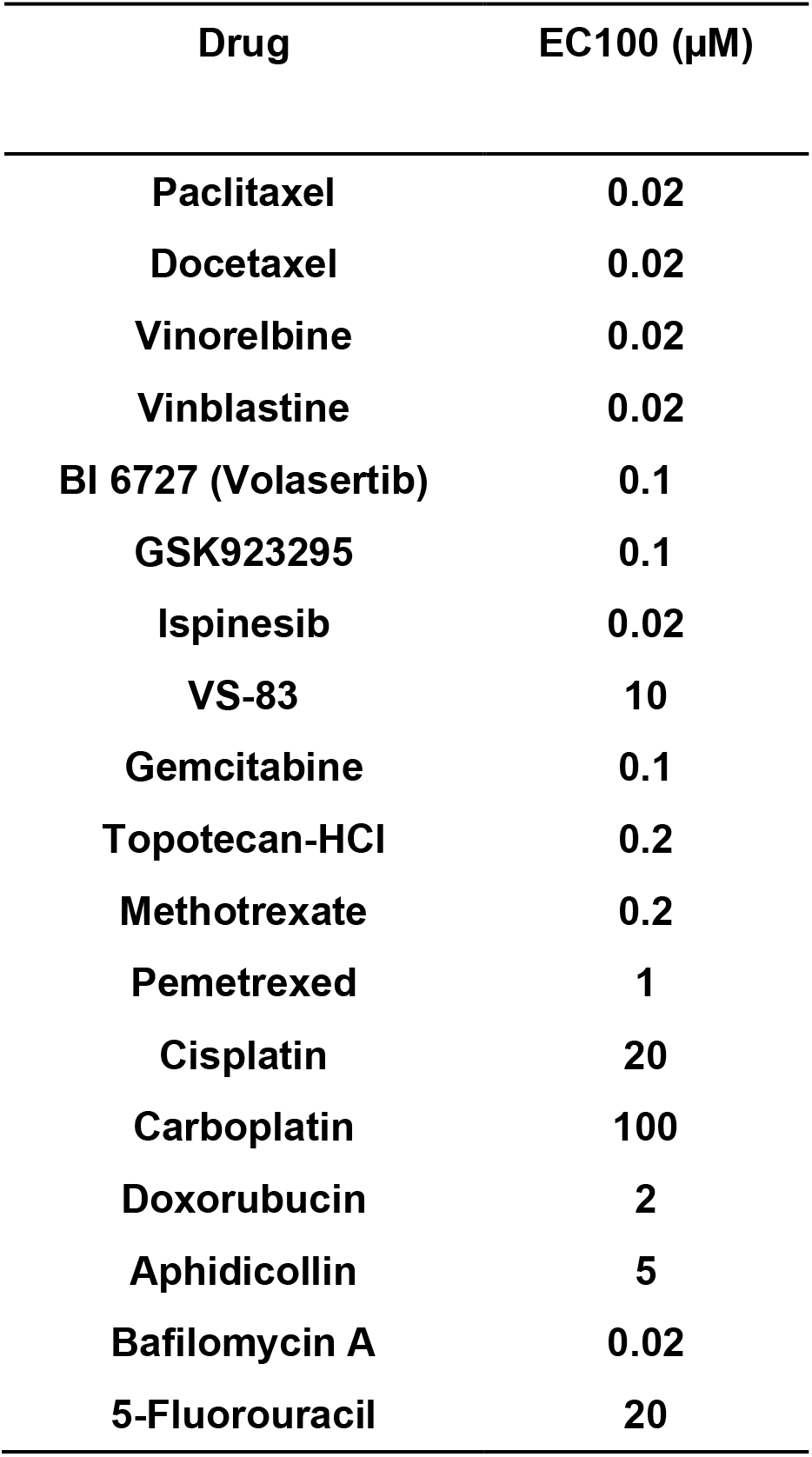
EC100 values of all used FDA-approved chemotherapy agents for NSCLCs or cytotoxic poisons in NCI-H1693.

